# Therapeutic vulnerability to ATR inhibition in concurrent *NF1* and *ATRX-*deficient high-grade solid tumors

**DOI:** 10.1101/2022.02.03.479021

**Authors:** Ming Yuan, Charles G. Eberhart, Christine A. Pratilas, Jaishri O. Blakeley, Christine Davis, Marija Stojanova, Karlyne Reilly, Alan K. Meeker, Christopher M. Heaphy, Fausto J. Rodriguez

## Abstract

Subsets of Neurofibromatosis Type 1 (NF1)-associated solid tumors have been shown to display high frequencies of ATRX mutations and presence of alternative lengthening of telomeres (ALT). In addition, a potential biologic vulnerability of ALT-positive cancer cells is ATR inhibition, a regulator of homologous recombination. We studied the phenotype of combined NF1 and ATRX deficiency in malignant solid tumors. For these experiments, cell lines derived from NF1-deficient sporadic glioblastomas (U251, SF188), a NF1-associated ATRX mutant glioblastoma cell line (JHH-NF1-GBM1), a NF1-derived sarcoma cell line (JHH-CRC65), and two NF1-deficient MPNST cell lines (ST88-14, NF90.8) were utilized. Cancer cells were treated with ATR inhibitors (AZD6738 and VE-822), in vitro, with or without a MEK inhibitor (AZD6244, selumetinib) or temozolomide. In contrast to the glioma cell line SF188, combined ATRX knockout (KO) and TERC KO led to ALT-like properties and sensitized U251 glioma cells to ATR inhibition (AZD6738 and VE-822) in vitro and in vivo. In addition, ATR inhibitors sensitized U251 cells to temozolomide, but not MEK inhibition (AZD6244), irrespective of ATRX level manipulation; whereas, the JHH-NF1-GBM1 cell line (ATRX loss/ALT-positive) demonstrated sensitivity to ATR inhibition (AZD6738), but not temozolomide. Similar effects were noted using the MPNST cell line NF90.8 after combined ATRX knockdown and TERC KO; however, not in the MPNST cell line ST88-14. Taken together, our study supports the feasibility of targeting the ATR pathway in subsets of NF1-deficient and associated tumors. Tumors with pre-existing ALT, or that subsequently develop ALT after ATRX downregulation, are particularly vulnerable to this therapeutic approach.

## Introduction

Neurofibromatosis type 1 (NF1) is an inherited tumor predisposition syndrome, and individuals affected with NF1 are prone to develop tumors of the central nervous system (CNS) and peripheral nervous system. Rarely, these individuals may develop other solid neuroendocrine and mesenchymal tumors, some of which may behave in a malignant fashion (1). The most common primary CNS tumors in these patients are pilocytic astrocytomas (PA) which have a predilection for the optic pathways, especially in children. It is also known that the full spectrum of glial neoplasia may develop in these patients, including low- and high-grade astrocytomas (2). Specific drivers of sporadic high-grade astrocytomas have been elucidated in the past years through genome-wide sequencing (3), and similar techniques have been applied to study NF1-associated astrocytomas, identifying genetic alterations in *TP53*, *CDKN2A*, and *ATRX* (4).

Telomeres are composed of a repetitive DNA sequence (TTAGGG) bound by a shelterin protein complex that protects the ends of linear chromosomes to maintain genomic integrity. However, in normal cells, telomeres progressively shorten with each cell division. Critically short telomeres can lead to breakage-fusion-bridge cycles that may lead to the accumulation of catastrophic global genomic damage or cellular senescence. To ensure unlimited replication potential, cancer cells employ two major, largely mutually exclusive, mechanisms of telomere maintenance. The predominant mechanism is expression of the reverse transcriptase, telomerase, which is encoded by an RNA component (*TERC*) and the rate limiting catalytic subunit (*TERT*). In cancer, telomerase up-regulation is frequently mediated through activating mutations (5), amplifications, structural variants, and promoter methylation (6) in *TERT*.However, a subset of cancers maintain their telomere lengths independent of telomerase, using the *<u>A</u>lternative <u>L</u>engthening of <u>T</u>elomeres* (ALT) (7). ALT results from an aberrant homologous recombination-based process mediated by homology-directed repair that leads to the presence of unique molecular features in ALT-positive cancers, including the presence of ultra-long telomeres, dramatic cell-to-cell telomere heterogeneity (assayable in cell and tissue samples via telomere-specific FISH), and the presence of single stranded extrachromosomal circles containing the C-rich telomere repeat sequence (C-circles; assayable by rolling circle amplification). Variations in telomere length have emerged as a prognostic factor in a variety of tumor types (8,9). Our group has reported a high frequency of ALT in high-grade astrocytomas developing in individuals with NF1-syndrome, as well as a small subset of MPNST, although usually not in their benign counterparts (10).

Prior studies have linked alterations in the *alpha thalassemia/mental retardation syndrome X-linked* (*ATRX*) or *death domain-associated protein* (*DAXX*) genes with ALT in a subset of cancers (11). Loss of ATRX function leads to abnormal methylation and gene expression patterns, as well as chromosome mis-segregation. In the nucleus, ATRX cooperates with the molecular chaperone DAXX to incorporate the H3.3 histone variant in heterochromatic regions, including at telomeres. *ATRX* mutations and ALT are associated with specific molecular subgroups of sporadic brain tumors (4,10). In NF1-associated tumors, particularly astrocytomas, *ATRX* mutations and ALT are associated with specific tumor groups such as the recently recognized WHO tumor type high-grade astrocytoma with piloid features (12,13).

Prior studies have documented that ATRX loss and ALT may represent a therapeutic vulnerability (14,15) and that these cells are sensitive to ATR inhibition in sporadic tumorigenesis (16). Additionally, ATRX inactivation promotes DNA damage and cellular death, which may synergize with specific therapeutic approaches. Thus, we hypothesized that the *ATRX* mutations and telomere alterations that occur in distinctive subsets of NF1-associated tumors, particularly those with an aggressive phenotype, represent a potential vulnerability that can be therapeutically targeted.

## Methods

### Cell Lines

Human tumor specimens were collected at Johns Hopkins Hospital with local Institutional Review Board approval, and written informed consent was obtained from patients or their parents. Tissues were minced and digested with either papain dissociation system Grand Island, New York), then filtered through a 70-μm Falcon cell strainer (ThermoFisher Scientific). JHH-NF1-GBM1 was grown in DMEM/F12 containing 1x B27 supplement (ThermoFisher Scientific), 20ng/ml EGF (PeproTech), 20ng/ml FGF-b (PeproTech) and 5μg/ml Heparin (Millipore SIGMA). Conditionally reprogramming culture (CRC) cell lines (JHH-NF1-PA1 and JHH-CRC65) were grown under the conditions described previously (17). 50% F medium (25% F-12, 75% DMEM supplied with 10% FBS and 5μg/ml Insulin), 50% 3T3 condition medium is supplied with 25 ng/ml Hydrocortisone (Millipore Sigma), 0.1 nmol/l Cholera toxin (Millipore SIGMA), 10 ng/ml EGF (PeproTech) and 5 μM ROCK inhibitor Y-27632 (Selleckchem). U251 was purchased from the American Type Culture Collection (ATCC) and SF188 was kindly provided by Dr. Chris Jones (Institute of Cancer Research, Sutton, UK). U251 ATRX^**-/-M**^, U251 ATRX^**-/-2.02**^, SF188 ATRX^**-/-F**^ and SF188 ATRX^**-/-N**^ were utilized as previously described (18). U251 and SF188 cells were grown in DMEM/F12 supplemented with 10% FBS (ThermoFisher Scientific). NF90.8 and ST88-14 were maintained in RPMI supplemented with 10% FBS (ThermoFisher Scientific). Murine glioma cell lines 130G#3, 158D#8, 1491-9 and 1861-10 were maintained in DMEM supplemented with 10% FBS (ThermoFisher Scientific) (19). All cells were cultured in a humidified 37°C incubator with 5% CO2. Human Schwann cells were purchased from ScienCell and maintained in Schwann cell media. Cell lines were routinely tested for mycoplasma and human cell line identities were confirmed by short tandem repeat (STR) profiling (Johns Hopkins University Genetic Resource Core Facility).

### Quantitative real-time polymerase chain reaction (qRT-PCR)

Total RNA was isolated from cultured cells using the RNeasy mini kit (QIAGEN), and cDNAs were produced using QuantiTect reverse transcription kit (QIAGEN). qRT-PCR was performed using PowerUp SYBR Green Master Mix (ThermoFisher Scientific). Primer sequences were ATRX: forward 5’-CAATCACAGAAGCCGACAAG -3’, reverse 5’-GTCATGAAGCTTCTGCACCA -3’; CDKN1A forward 5’-AGTCAGTTCCTTGTGGAGCC -3’, reverse 5’-CATGGGTTCTGACGGACAT -3’; CDKN1B forward 5’-AAGAAGCCTGGCCTCAGAAG-3’, reverse 5’-TTCATCAAGCAGTGATGTATCTGA-3’; CDKN2A forward 5’-GTTACGGTCGGAGGCCG -3’, reverse 5’-GTGAGAGTGGCGGGGTC -3’; PARP1 forward 5’-GATGGGTTCTCTGAGCTTCG -3’, reverse 5’-TCTGCCTTGCTACCAATTCC -3’; TERC forward 5’-CCCATTCATTTTGGCCGACTT -3’, reverse 5’-GGCCGCTCCCTTTATAAGC -3’; HPRT1 forward 5’-GTTATGGCGACCCGCAG-3’, reverse 5’-ACCCTTTCCAAATCCTCAGC-3’. HPRT1 was used as the endogenous control. The relative fold changing was calculated based on the formula R=2^-(ΔCt sample-ΔCt control)^. Quantification qPCR assay for telomerase activity was performed according to the manufacturer’s instructions (#8928, ScienCell).

### Gene knockdown and knockout of ATRX or TERC

The CRISPR cas9 nickase system was used to generate inactivating mutations in either the *ATRX* or *TERC* genes, as previously reported (20,21). Guide RNAs (gRNAs) targeting TERC were obtained from Abmgood. Short hairpin RNA target human ATRX (sh11 and sh90), mouse ATRX (sh1 and sh3) and vector control pLKO.1 were obtained from Millipore Sigma. To produce lentiviruses, 293T cells were transfected with shRNA or gRNA plasmid and VSVG packaging plasmids mixture using lipofectamine 2000 (ThermoFisher Scientific). Lentiviral supernatants were collected at 48-72 h later and kept frozen at −80°C until needed. Cells infected with virus were selected with 1-2 μg/ml of puromycin (MilliporeSigma), 2-4 μg/ml of Blastidin (ThermoFisher Scientific) and 100-400 μg/ml of G418 (ThermoFisher Scientific) for 7 days to generate stable cell lines. ATRX knockdown was confirmed by western blotting and *TERC* knockout (KO) was confirmed by qPCR.

### ALT validation

ALT was assessed with previously established methods (22–24). Ultra-bright telomeric foci were evaluated by telomere-specific FISH. Telomeric extra-chromosomal circles (e.g. C-circles) were detected using immunoblotting after a processive phi29 polymerase to amplify C-circle DNA. A DIG-conjugated probe containing the C-rich telomere repeat sequence specifically targeted the polymerase amplified signal. The known ALT-positive osteosarcoma cell line U2OS served as a positive control.

### Cell growth assessment

To assess effects on cell growth, the CellTiter-Blue assay was used (Promega). In brief, 1000 to 5000 cells were plated in triplicate in 96-well plates. 20-30 microliters of the CellTiter-Blue reagent was added per well in 96-well plates and incubated for 1-4 h at 37°C in 5% CO2. For drug treatments, cells in 96-well plates were cultured with various drug concentrations of AZD6244, AZD6738, VE-822, or temozolomide (Selleckchem). Vehicle (Dimethyl sulfoxide)-treated cells were used as controls and the cell survival fraction was calculated as percentage of control cells. Fluorescence (560 nmEx/590 nmEm) was measured using TECAN plate reader. Additionally, apoptosis assays were performed using Muse Annexin V & Dead Cell reagent (MilliporeSigma) and Bromodeoxyuridine (BrdU) incorporation assays were performed as previous described (25). Data were acquired using Muse flow cytometer (Millipore) and analyzed with FlowJo software.

### Western blotting

Cells were lysed in RIPA lysis buffer supplemented with protease inhibitors (MilliporeSigma). Primary antibodies used for western blots were: NF1 (A300-140A, 1:1000, Bethyl Laboratories), ATRX (#10321, 1:500, Cell Signaling Technology), α-tubulin (#3873, 1:5000, Cell Signaling Technology), β-actin (sc-47778, 1:5000, Santa Cruz Biotechnology), pERK1/2 (#4370, 1:1000, Cell Signaling Technology), Erk1/2 (#9102, 1:1000, Cell Signaling Technology),γH2AX (#9718, 1:1000, Cell Signaling Technology). Secondary antibodies used for western blots were antimouse IgG HRP-linked (#7076, 1:5000, Cell Signaling Technology) and anti-rabbit IgG HRP-linked (#7074, 1:5000, Cell Signaling Technology).

### MPNST xenograft

For *in vivo* experiments, 1×10^6^ cells were sciatic nerve transplanted in Nude mice (Charles River). For bioluminescence imaging, cells were labelled with the lentiviral based reporter co-expressing RFP and luciferase (SBI). Animals were closely monitored for tumor growth, and euthanized when tumor size was over 1000 mm^3^

### *In vivo* drug testing

For *in vivo* experiments, 5×10^5^ glioma cells were orthotopically transplanted (Stereotaxic coordinate: X(AP)=1.0 mm, Y(ML)=2.0 mm, Z(DV)=-3 mm) in Nude mice (Charles River). For bioluminescence imaging, cells were labelled with the lentiviral based reporter co-expressing RFP and luciferase (SBI). Oral drugs were delivered to xenografted mice once daily by gavage from d2 to d16. Animals were closely monitored for tumor growth, and euthanized when neurologic signs of disease develop. In addition, subsets were sacrificed at appropriate intervals and tissue sections microscopically examined for early evidence of tumor formation. Methods for the analysis of tumor xenograft for morphology, size, proliferation and differentiation are previously reported (26–28).

### Statistical Analyses

For cell culture and functional assays, data were presented as mean ± standard deviation with p<0.05 considered statistically significant. All experiments were performed in at least three biological replicates and data analyzed with a two-tailed Student’s t-test or ANOVA as appropriate. Survival was assessed with Kaplan-Meier curves and statistical analyses made using standard software and statistical packages (GraphPad).

## Results

### ATRX loss in the context of diminished telomerase activity facilitates the development of ALT-associated hallmarks

To identify the biologic relevance of ATRX loss in NF1-associated gliomagenesis, we studied four previously characterized *Nf1^+/−^Trp53^+/−^* murine glioma lines (130G#3, 158D#8, 1491-9, 1861-10) representing all of the diffuse glioma grades, 2-4 (19). Compared to NIH-3T3 cells, these cell lines all underexpress *Nf1* while demonstrating variable *Atrx* expression, with Atrx loss in two cell lines, and preserved Atrx expression in two cell lines (**Supplementary Figure 1**).

Since ALT can develop in the presence of diminished telomerase activity (29), we tested the effects of *Atrx* knockdown (cell lines 130G#3 and 158D#8) in the presence of the telomerase inhibitor, BIBR1532. This inhibitor decreases telomerase activity by binding to the active site of TERT, thereby downregulating TERT expression (30,31). With the known caveat that murine telomere lengths are significantly longer and display stronger telomeric FISH signals compared to human cancer cells, we identified increased telomere FISH signal brightness, with features reminiscent of ALT, after prolonged Atrx knockdown (55 days) and concurrent Tert inhibition (39 days) (**Figure 1**).

**Figure 1.**
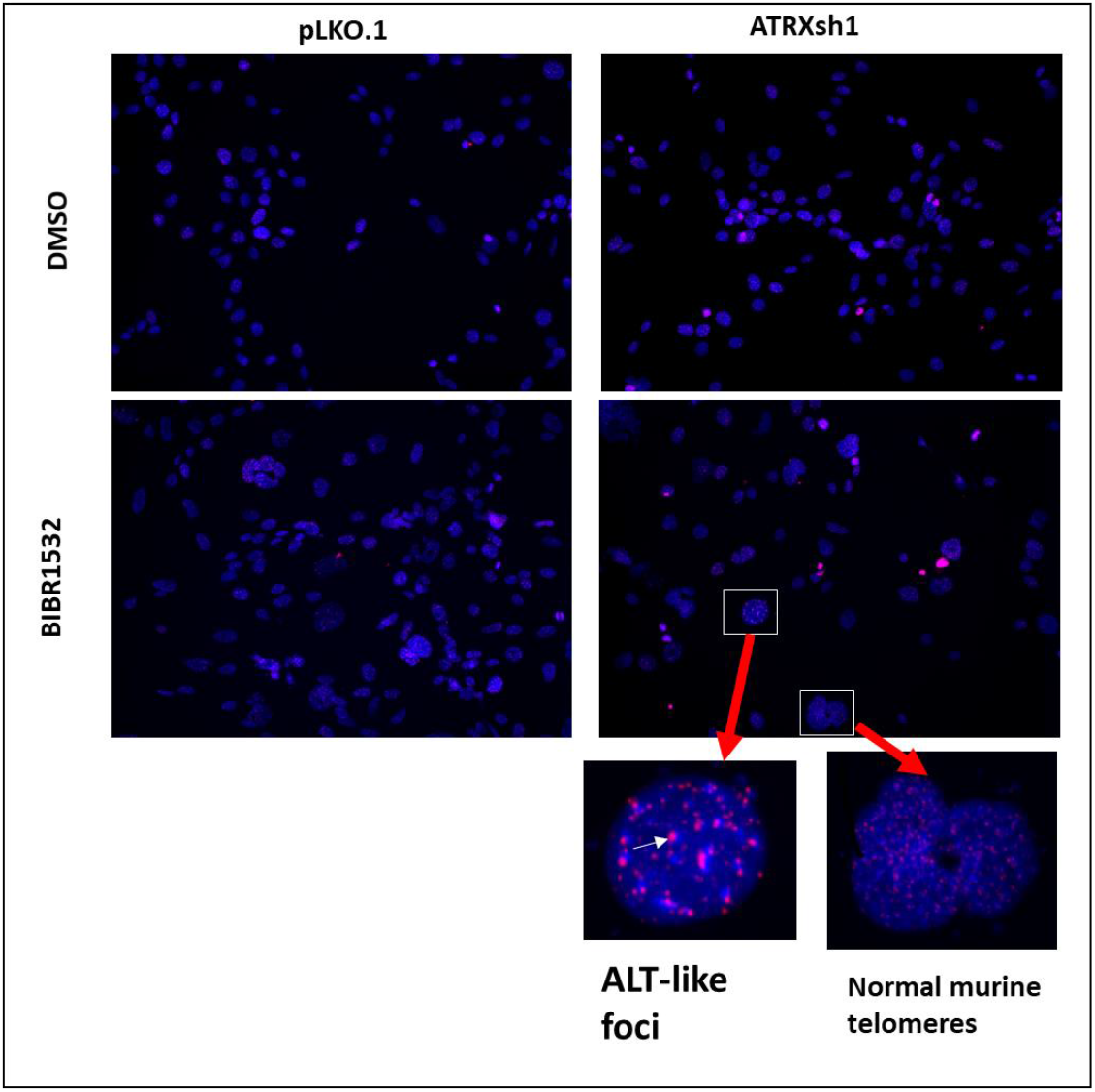
Telomere-specific FISH staining in *Nft^+/−^Tp53^+/−^* murine glioma cell line 130G#3. Development of ultrabright telomere foci, indicative of ALT, after *Atrx* knockdown and treatment with the telomerase inhibitor, BIBR1532 for 39 days.

### *ATRX* loss decreases cell growth in NF1-deficient human glioma lines, but with no effect in MPNST lines

Using the *Conditionally Reprogramming Culture* (CRC) technique (32), we previously developed a human pilocytic astrocytoma cell line derived from a patient with NF1 (17). *ATRX* depletion through shRNA resulted in decreased growth and increased apoptosis (**Figure 2**). Since ATRX loss and ALT activation are largely limited to high-grade neoplasms in the context of NF1 loss or inherited NF1 syndrome (10), we evaluated cell lines derived from high-grade neoplasms with *NF1* loss, including two sporadic glioblastoma cell lines with *NF1* inactivation (U251, SF188), a NF1-associated glioblastoma (JHH-NF1-GBM1), two MPNST cell lines (NF90.8, ST88-14) and a NF1-associated sarcoma line (JHH-CRC65). These cell lines displayed NF1 protein loss, preserved DAXX protein expression, but demonstrated variable ATRX protein levels. Notably, ATRX protein expression was higher in the sporadic glioma lines (U251, SF188), while completely absent in the NF1-associated glioblastoma line JHH-NF1-GBM1 (**Supplementary Figure 2**).

**Figure 2.**
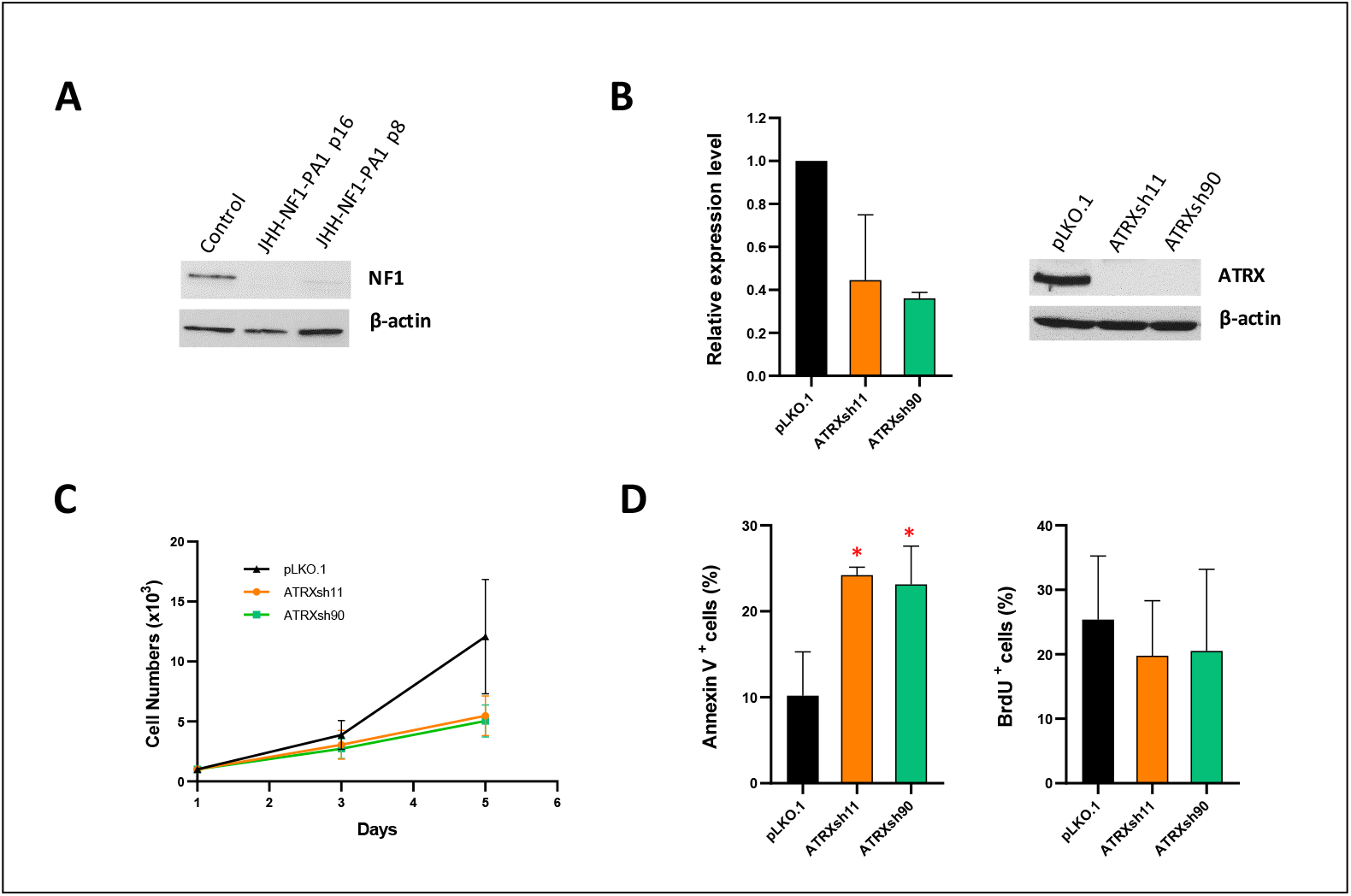
ATRX knockdown in the human glioma cell line JHH-NF1-PA1. NF1 pediatric pilocytic astrocytoma cell line JHH-NF1-PA1 has pronounced NF1 loss in culture (A). Successful *ATRX* knockdown (mRNA left, protein right) using shRNA hairpins (B). ATRX loss leads to decreased growth of JHH-NF1-PA1 (C), primarily through increased apoptosis (D).

ATRX knockdown (via shRNA) or KO (via CRISPR) resulted in decreased growth in the high-grade glioma cell lines (U251, SF188), with increased apoptosis more evident in U251 *in vitro* (**Figure 3A-B**). Reduced growth of U251 after ATRX KO was also evident *in vivo* using orthotopic intracranial xenografts upon histological examination (**Supplementary Figure 3**), leading to a prolonged survival (**Figure 3C**). Functional loss of ATRX resulted in ALT in U251, but not in SF188 (18), and ALT features were present in the NF1-glioblastoma line JHH-NF1-GBM1 as demonstrated by the c-circle assay (**Supplementary Figure 4**) and telomere-specific FISH.

**Figure 3.**
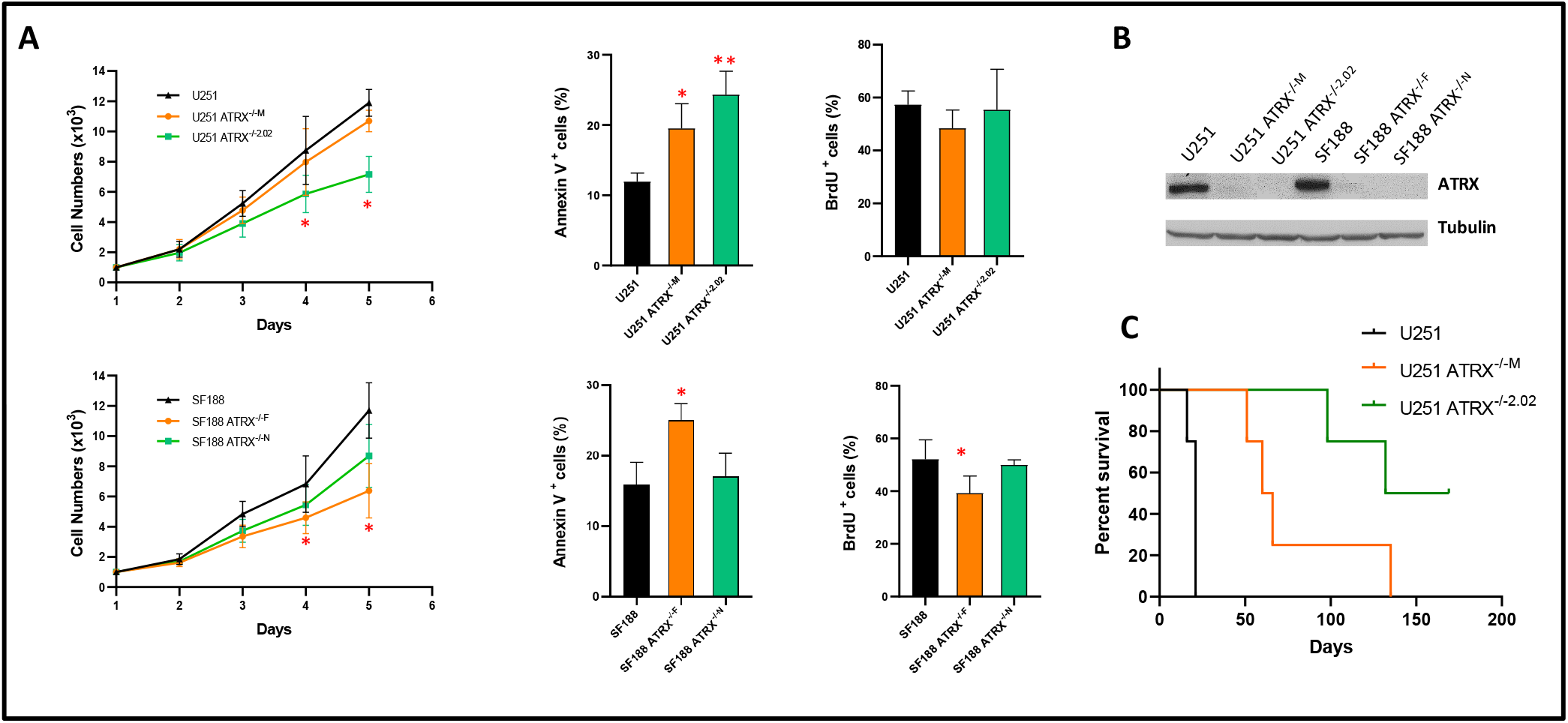
ATRX knockout in NF1-deficient glioblastoma cell lines U251 and SF188. Decreased cell growth, secondary to apoptosis, was noted *in vitro* (A). Successful ATRX knockout in both glioma lines at the protein level (B). ATRX knockout in U251 resulted in decreased tumor growth in orthotopic nude mouse xenografts (C).

Next, we studied the effects of ATRX loss in NF1-deficient MPNST and sarcoma cell lines. In contrast to the NF1-deficient glioma lines, and despite successful *ATRX* knockdown, ATRX loss had no effect on growth *in vitro* (NF90-8, ST88-14, JHH-CRC65) or *in vivo* (NF90-8) (**Supplementary Figures 5-7**). Telomere-specific FISH staining of these cells showed rare ultrabright, ALT-like foci (**Figure 4**). In contrast, *ATRX* knockdown resulted in decreased growth of a non-neoplastic Schwann cell line with increased levels of senescence marker p21 (CDKN1A) (**Supplementary Figure 8**).

**Figure 4.**
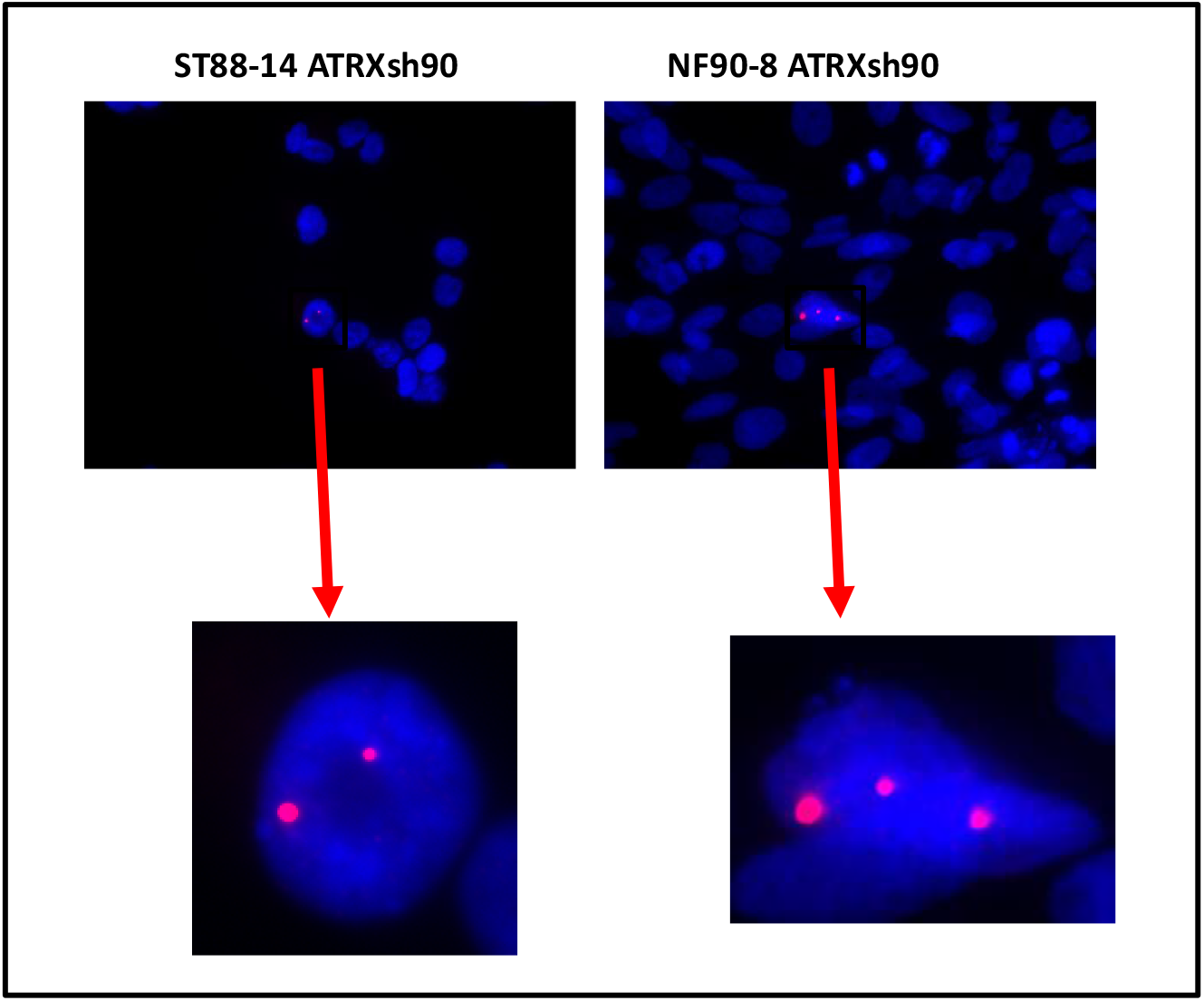
ATRX knockdown resulted in rare abnormal telomere FISH signals. Rare ultrabright telomeric foci were detected in the MPNST cell lines ST88-14 and NF90.8 after *ATRX* knockdown.

### ATRX deficiency sensitizes cells to ATR inhibitors and temozolomide

To test the feasibility of ATR inhibition in the treatment of high-grade neoplasms, we evaluated *NF1-* and *ATRX*-deficient cancer cells. The glioma cell line U251, which develops ALT after ATRX KO (18), was more sensitive to two different ATR inhibitors (AZD6738 and VE-822) in the absence of ATRX, in contrast to SF188 cell line which remains ALT-negative (**Figure 5**). In addition, ATRX KO also sensitized U251 to MEK inhibition (AZD6244) (**Figure 5**). Next, we tested the ATR inhibitor, AZD6738, in combination with temozolomide or MEK inhibition. The combination treatment of temozolomide and AZD6738 profoundly decreased growth in ATRX-deficient U251 cells (**Supplementary Figure 9**), while MEK inhibition (AZD6244) had no additional effect despite inhibiting MEK signaling as demonstrated by a decrease in pERK levels (**Supplementary Figure 10**). The NF1-associated glioblastoma cell line JHH-NF1-GBM1, which carries an *ATRX* mutation and exhibits ALT, grows under neurosphere conditions and intracranially as orthotopic xenografts, albeit slowly (**Figure 6A-B**). This cell line demonstrates sensitivity to ATR inhibition (AZD6738) *in vitro* while being resistant to temozolomide, a standard drug used in the treatment of high-grade glioma (**Figure 6C**). Similarly, *ATRX*knockdown and *TERC* KO in the MPNST cell line NF90.8 successfully suppressed telomerase activity (**Supplementary Figure 11**), and sensitized cells to ATR inhibition (AZD6738 and VE-822) (**Figure 7**). In contrast, *ATRX* knockdown in the ALT-negative, NF1-associated sarcoma line JHH-CRC65 did not increase sensitivity to either ATR inhibition or temozolomide treatment (**Supplementary Figure 12**).

**Figure 5.**
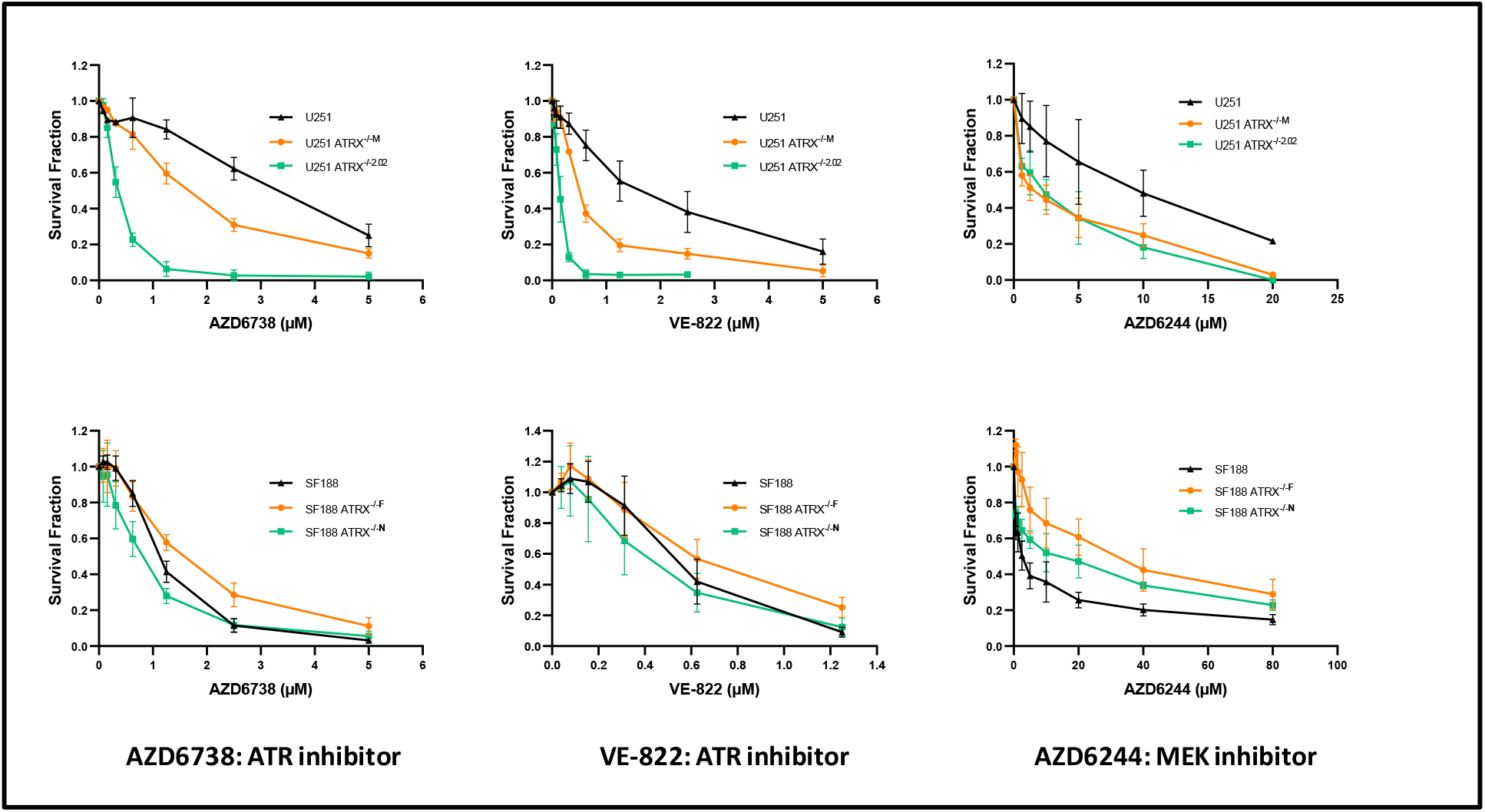
ALT-positive, ATRX-deficient U251 cells display an increased sensitivity to ATR inhibition. U251 and SF188 cells were treated with various doses of AZD6244, VE-822 or AZD6738 for 5 days, cell survival were normalized with vehicle control. AZD6738 and VE-822 decreased growth of U251 (top row), but not in cell line SF188 (ALT-) (bottom row). MEK inhibition (AZD6244, selumetinib) had a modest effect on growth in U251 as well.

**Figure 6.**
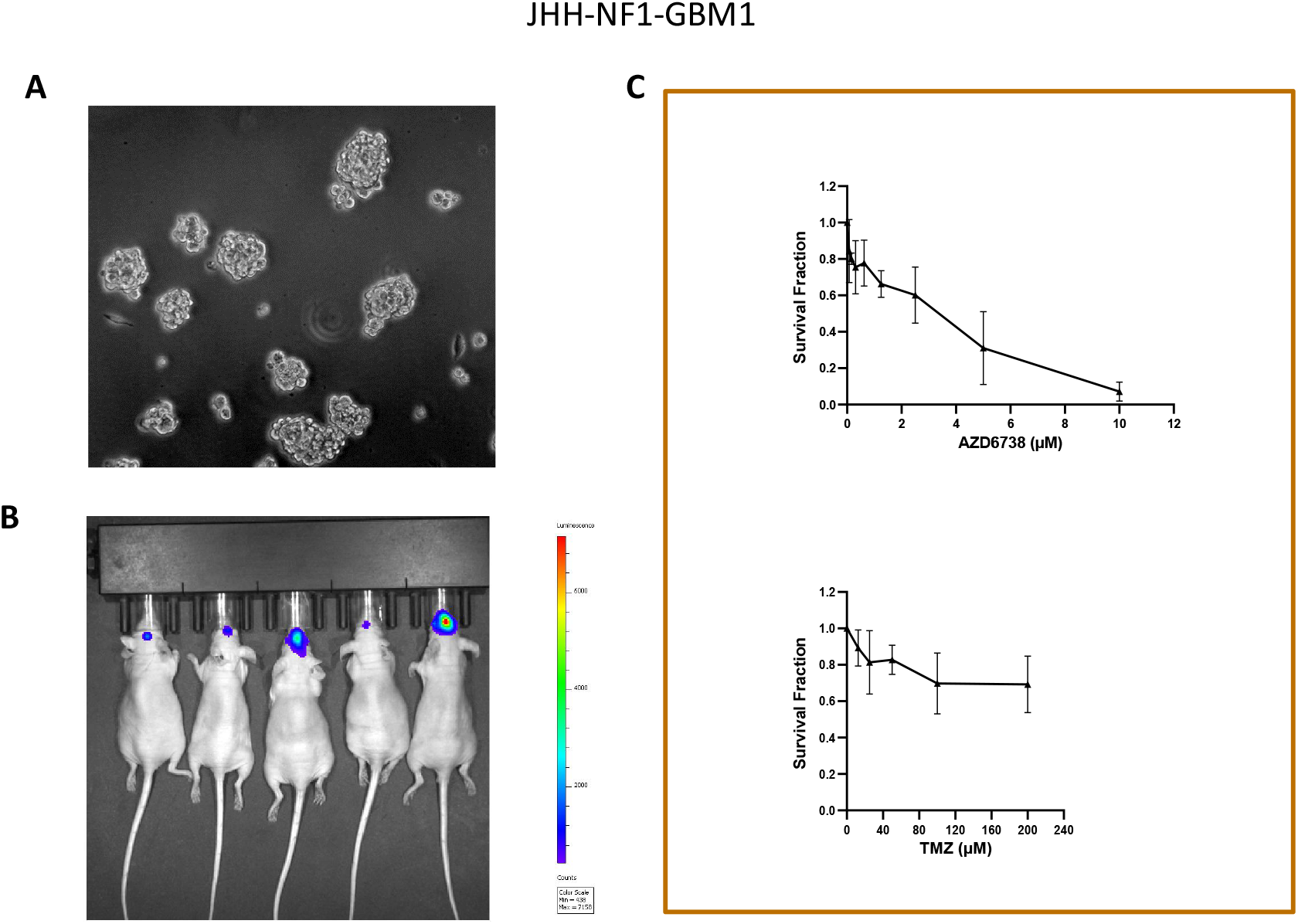
ATR inhibition impairs growth of the NF1-patient derived glioblastoma cell line JHH-NF1-GBM1 *in vitro*. JHH-NF1-GBM1 grown under neurosphere culture condition (A) and in orthotopic nude mouse xenografts (B). Decreased cell growth after treatment with the ATR inhibitor, AZD6738, was noted *in vitro*, although these cells remained relatively resistant to treatment with temozolomide.

**Figure 7.**
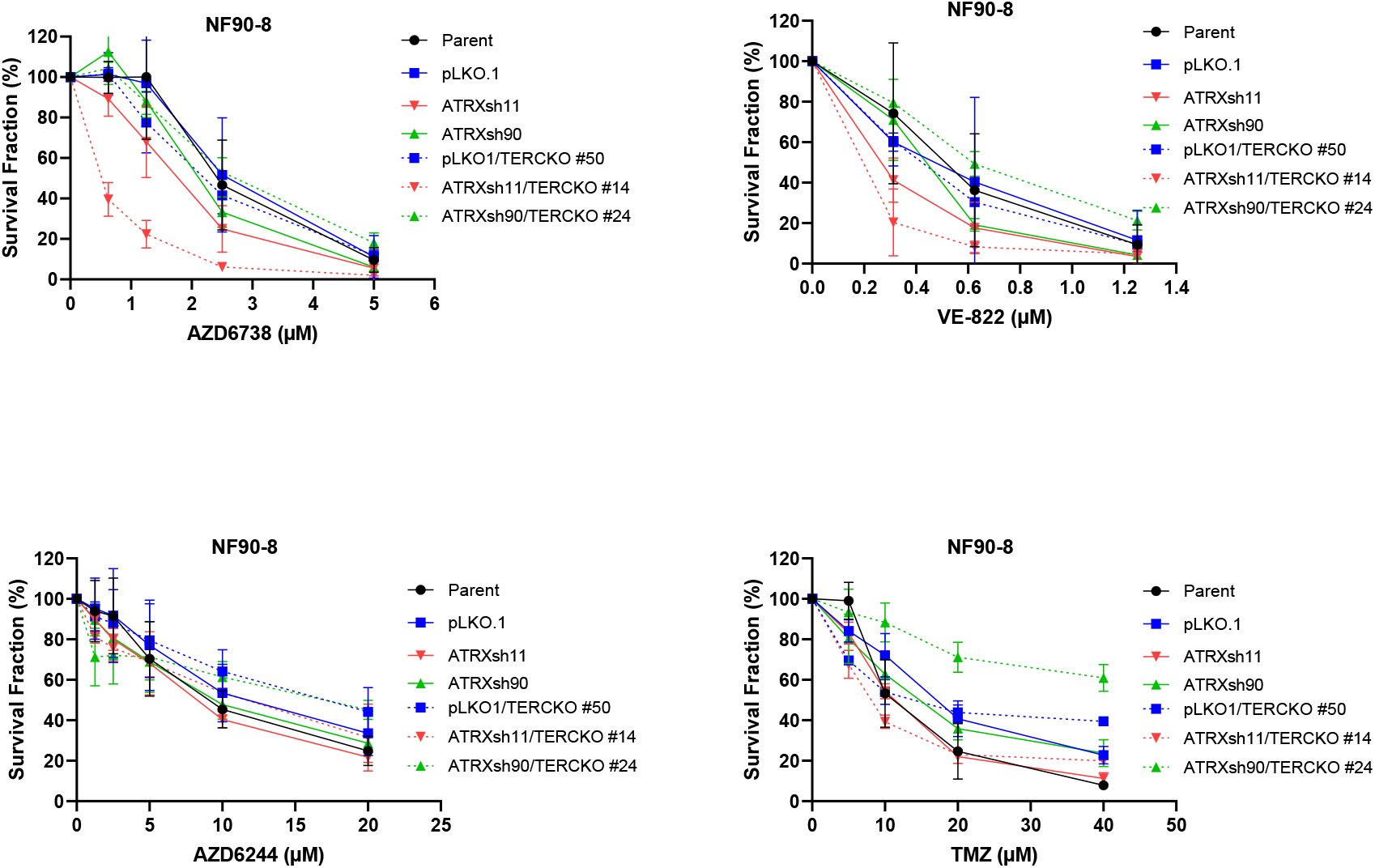
ATRX and TERC loss impairs growth of the MPNST cell line NF90-8 when treated with ATR inhibitors. MPNST cell clones with concurrent ATRX knockdown and TERC knockout are sensitive to ATR inhibitors AZD6738 and VE-822, but not to MEK inhibitor (AZD6244) or temozolomide.

Finally, we investigated the effect of ATR inhibition and temozolomide treatment on orthotopic glioma xenografts *in vivo*. Oral administration of the ATR inhibitor, AZD6738, decreased growth transiently in *ATRX*-deficient U251 xenografts, while the combination of temozolomide plus AZD6738 had a more pronounced and persistent inhibitory effect on tumor cell growth (**Figure 8**).

**Figure 8.**
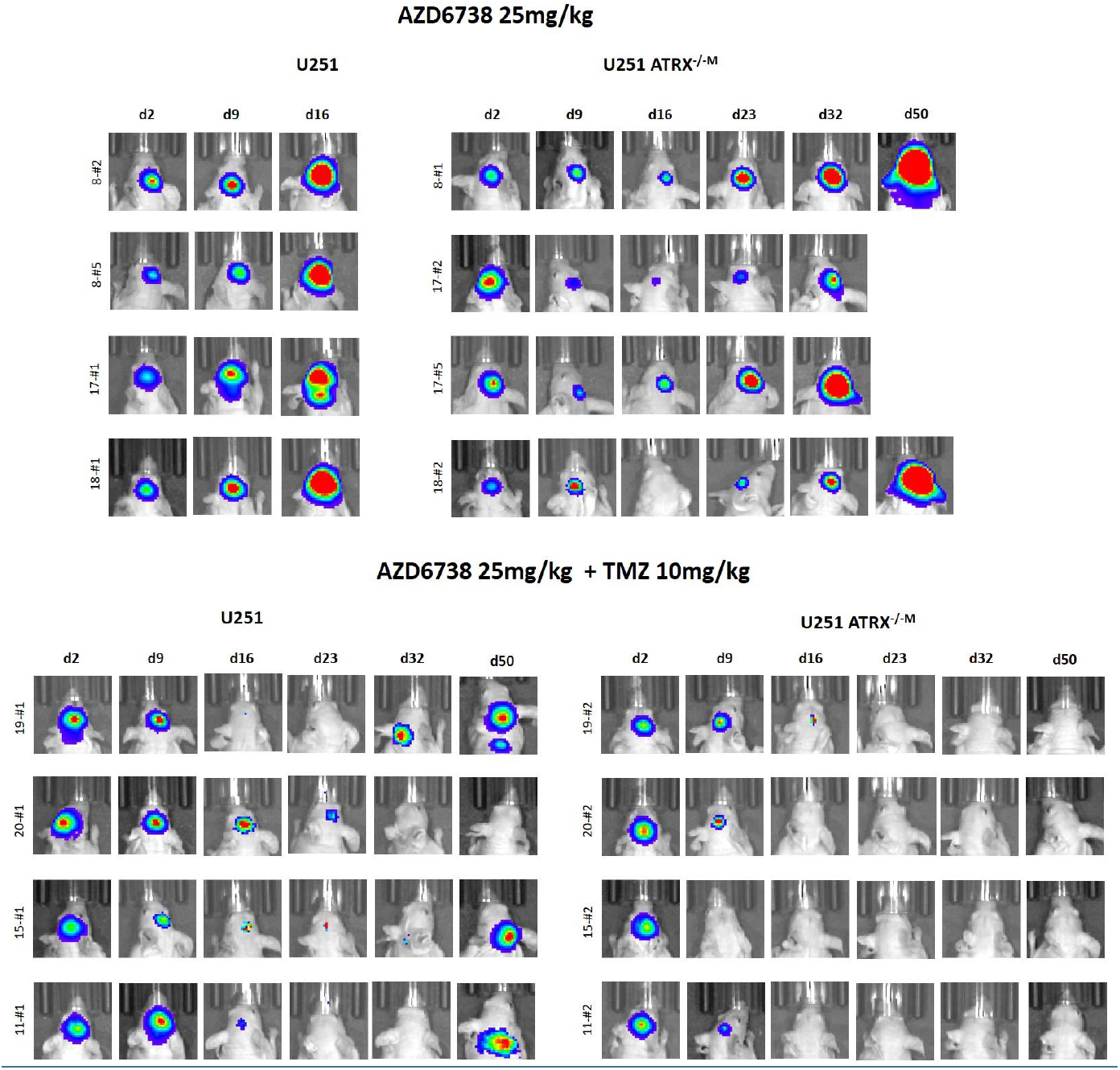
ATR inhibition transiently impairs growth in the NF1 glioma cell line U251 with ATRX loss. Orthotopic xenografts using the glioma cell line U251 demonstrate transient growth inhibition after treatment with the ATR inhibitor, AZD6738. Growth inhibition is more persistent when combined with temozolomide.

## Discussion

Inactivation of ATRX, or, less frequently, its associated protein DAXX, through deleterious mutations has been increasingly observed in subsets of cancers, and is consistently associated with ALT. There has been increasing interest in the biologic effect of ATRX, and particularly whether targeted therapies may be helpful for these tumors, which are often high-grade and difficult to treat. Functional loss of ATRX leads to G-quadruplex (G4) DNA secondary structures as a result of replication stress in glioma models (33). For example, *Atrx*-null neural progenitor cells are exquisitely sensitive to telomestatin, a compound that stabilizes G4 DNA secondary structures (34). Several groups have also reported on the efficacy of specific drugs for *ATRX*-mutant gliomas. In a mouse model developed using the *Sleeping Beauty* transposon system, Koschmann et al. demonstrated that Atrx deficiency leads to impaired non-homologous end joining, and sensitizes cells to compounds associated with the formation of double stranded DNA breaks (35). More recently, this group demonstrated that ATRX-deficient cells are associated with loss of Chk1 and reliant on ATM, thereby suggesting that ATM inhibition may sensitize these cells to radiation therapy (36). In addition, ATRX depletion in cells results in persistent association of telomeres with replication protein A, which in the presence of ATR inhibition, results in disruption of ALT and selective cell death (16). Several ATR inhibitors are in active phase I/II (37), including VX-970 and AZD6738, with AZD6738 having the added benefit of being orally bioavailable.

Despite the observation that ATRX-deficient tumors essentially always develop ALT, this is not consistently observed in model systems, suggesting that in certain cellular contexts, ATRX loss is not sufficient to induce ALT (18,21,38). Our experience demonstrates that attenuated telomerase activity (i.e. telomerase inhibition) is necessary, which may require prolonged or permanent TERT inactivation, and our data suggests this phenomenon to be cell-type dependent. The focus of our study is on functional loss of ATRX, since it is the predominant gene inactivated by mutations in solid tumors associated with ALT, particularly tumors arising from the central and peripheral nervous systems. It seems that indeed, ALT-positive cells may require the presence of specific DNA repair proteins, but the precise requirements vary by cell type (39).

In summary, ATRX loss and the development of ALT in NF1-deficient tumors represents a potential therapeutic vulnerability to ATR inhibition that may lead to much needed targeted treatments for solid tumors developing in patients with NF1 or in the sporadic setting. However, this vulnerability may be cancer type and/or cell line dependent, and therefore further studies are needed to identify the best biomarkers and/or drugs that will help manage this patient population.

## Acknowledgments

This work was supported in part by DOD W81XWH-18-1-0496 (FJR, CH, AM), Pilocytic/Pilomyxoid Fund, including Lauren’s First and Goal, and the Stick it to Brain Tumors Annual Women’s Ice Hockey Tournament (F.J.R, CGE.) and NIH grant P30 CA006973 to the Sidney Kimmel Comprehensive Cancer Center (PI: W. Nelson).

