## Supplemental Figures 1-12 for "Therapeutic vulnerability to ATR inhibition in concurrent *NF1* and *ATRX-*deficient high-grade solid tumors"

**Supplementary Figure 1. Atrx knockdown in *Nf1*$^{+/-}$*Tp53*$^{+/-}$ murine glioma lines (130G#3 and 158D#8) did not affect cell growth.** These cell lines demonstrate variable Atrx expression (A). Two lines with Atrx expression were used for knockdown experiments (B). There were no significant effects on cell growth (C).

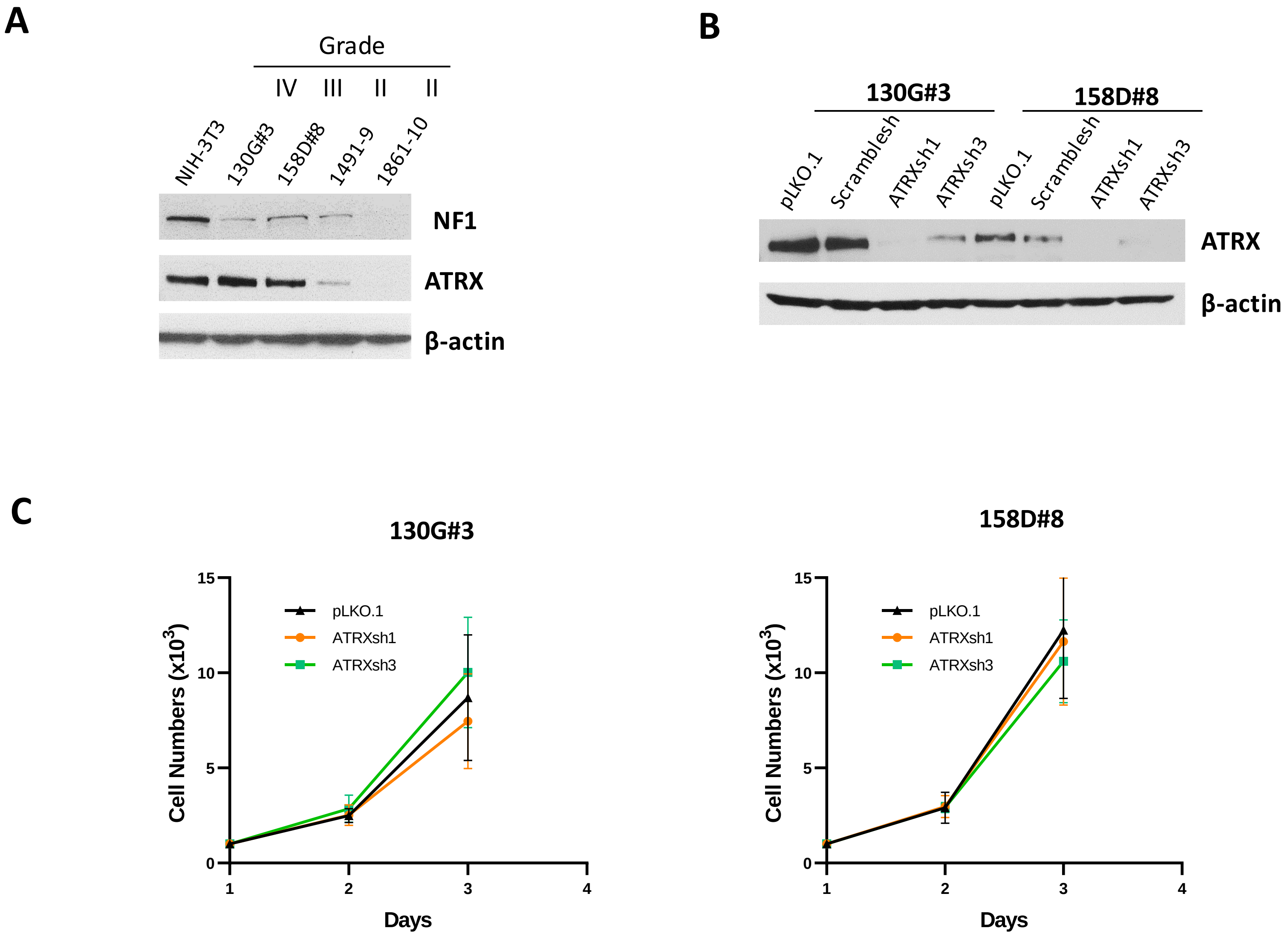

**Supplementary Figure 2. NF1-deficient malignant cell lines and ATRX expression.** Cell lines studied included sporadic glioma cell line U251 and SF188, NF1-patient derived cell line JHH-NF1-GBM1, NF1-patient derived sarcoma line JHH-CRC65, and MPNST cell lines NF90.8 and ST88-14. JHH-NF1-GBM1 was derived from a glioblastoma in a NF1 patient. In addition, there was complete ATRX protein loss and an *ATRX* mutation.

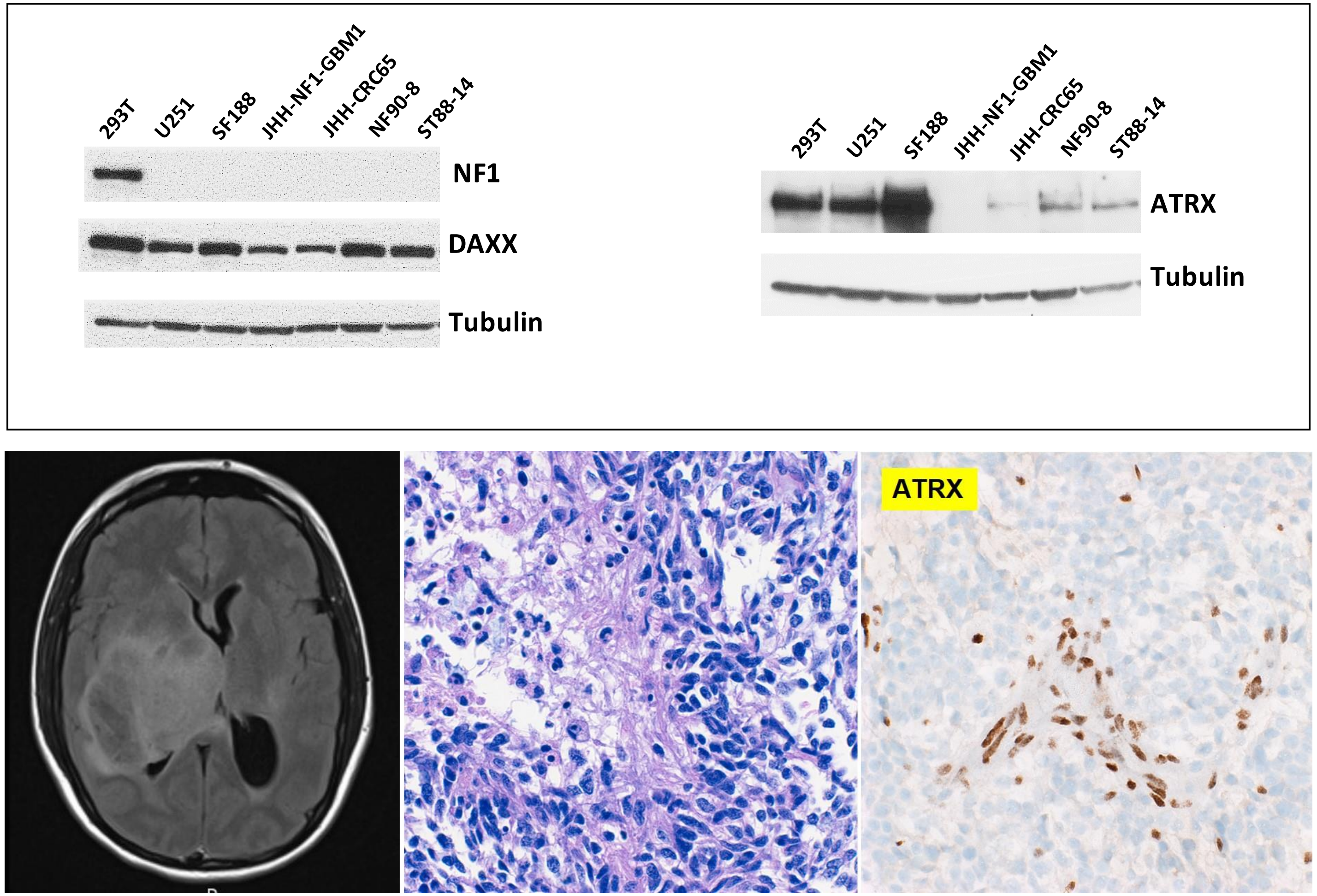

**Supplementary Figure 3. ATRX knockout in NF1-deficient high-grade glioma *in vivo*.** ATRX knockout resulted in less tumor burden after ATRX loss. Representative coronal sections of murine brains representing orthotopic xenografts of cell line U251 with and without ATRX knockout are presented.

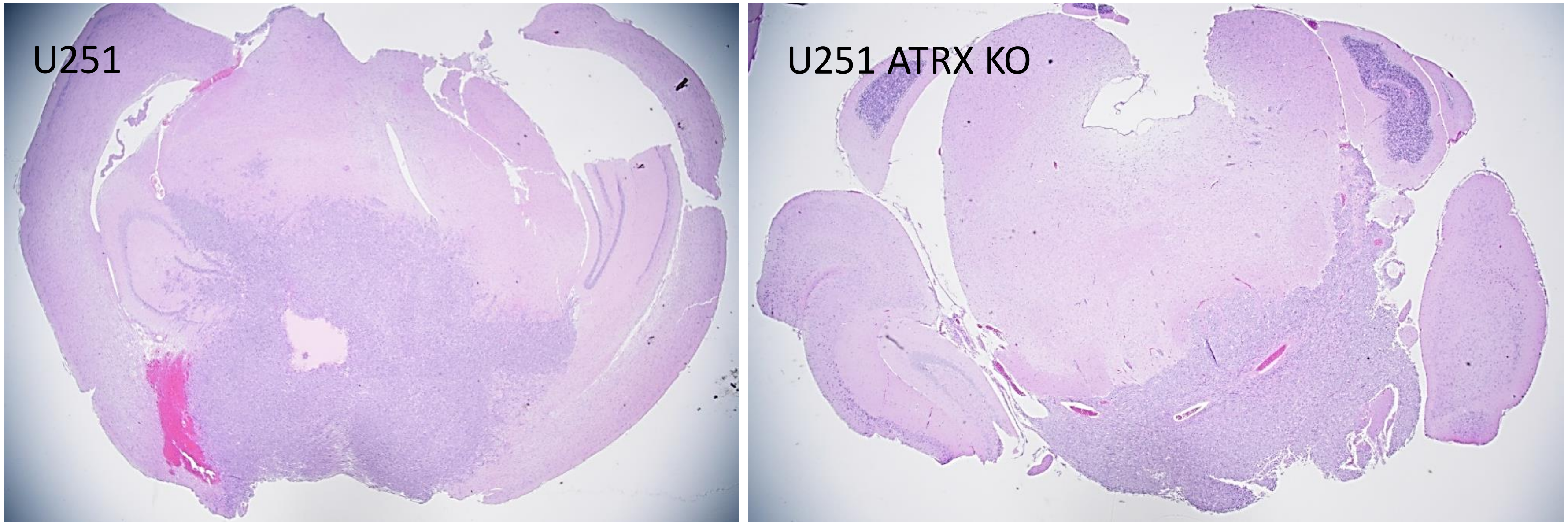

**Supplementary Figure 4. ATRX loss and ALT in NF1-deficient cell lines.** ATRX loss by knockout (U251) or preexisting mutation (JHH-NF1-GBM1) resulted in ALT in these two glioma cell lines as demonstrated by a c-circle assay. By contrast, NF1-sarcoma derived cell line JHH-CRC65 was ALT negative. Appropriate positive control (U2OS ALT-positive osteosarcoma cell line) and negative controls are shown at the top.

| | |
|---|---|
| 1 | U251 |
| 2 | U251/TERCKO#2 |
| 3 | U251/TERCKO#8 |
| 4 | U251 |
| 5 | U251 ATRX-/-M/TERCKO#13 |
| 6 | U251 $ATRX^{-/-M}$ /TERCKO#33 |
| 7 | U251 $ATRX^{-/-M}$ /TERCKO#42 |
| 8 | U251 $ATRX^{-/-2.02}$ |
| 9 | U251 $ATRX^{-/-2.02}$ /TERCKO#30 |
| 10 | U251 $ATRX^{-/-2.02}$ /TERCKO#51 |
| 11 | JHH-NF1-GBM1 |
| 12 | JHH-CRC65 |

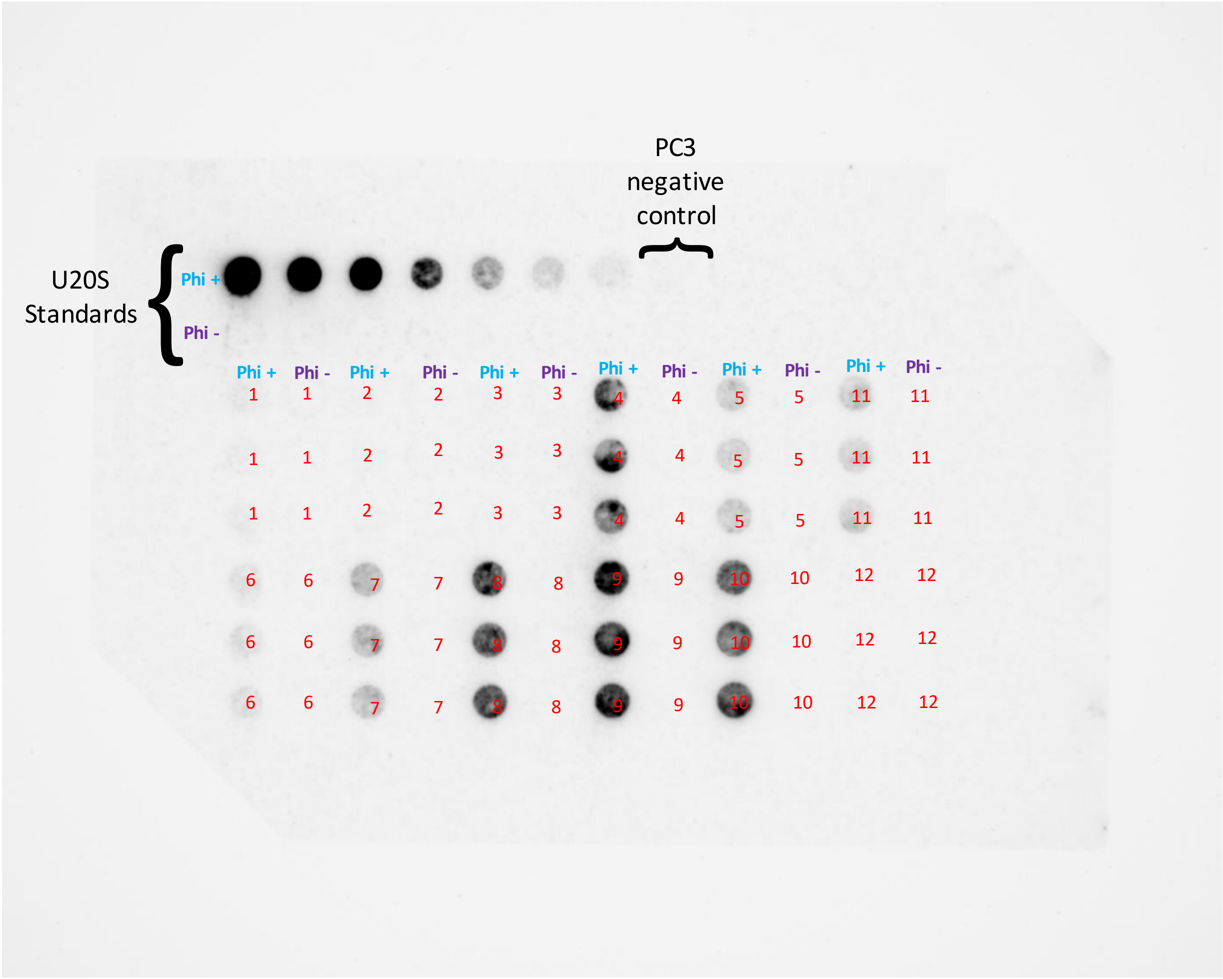

**Supplementary Figure 5. ATRX knockdown in NF90-8 has no effect on cell growth.** Successful ATRX knockdown in the MPNST cell line NF90-8 has no significant effect on cell growth *in vitro* (A) or *in vivo* (B). Illustrated xenografts were performed with perisciatic nerve injections.

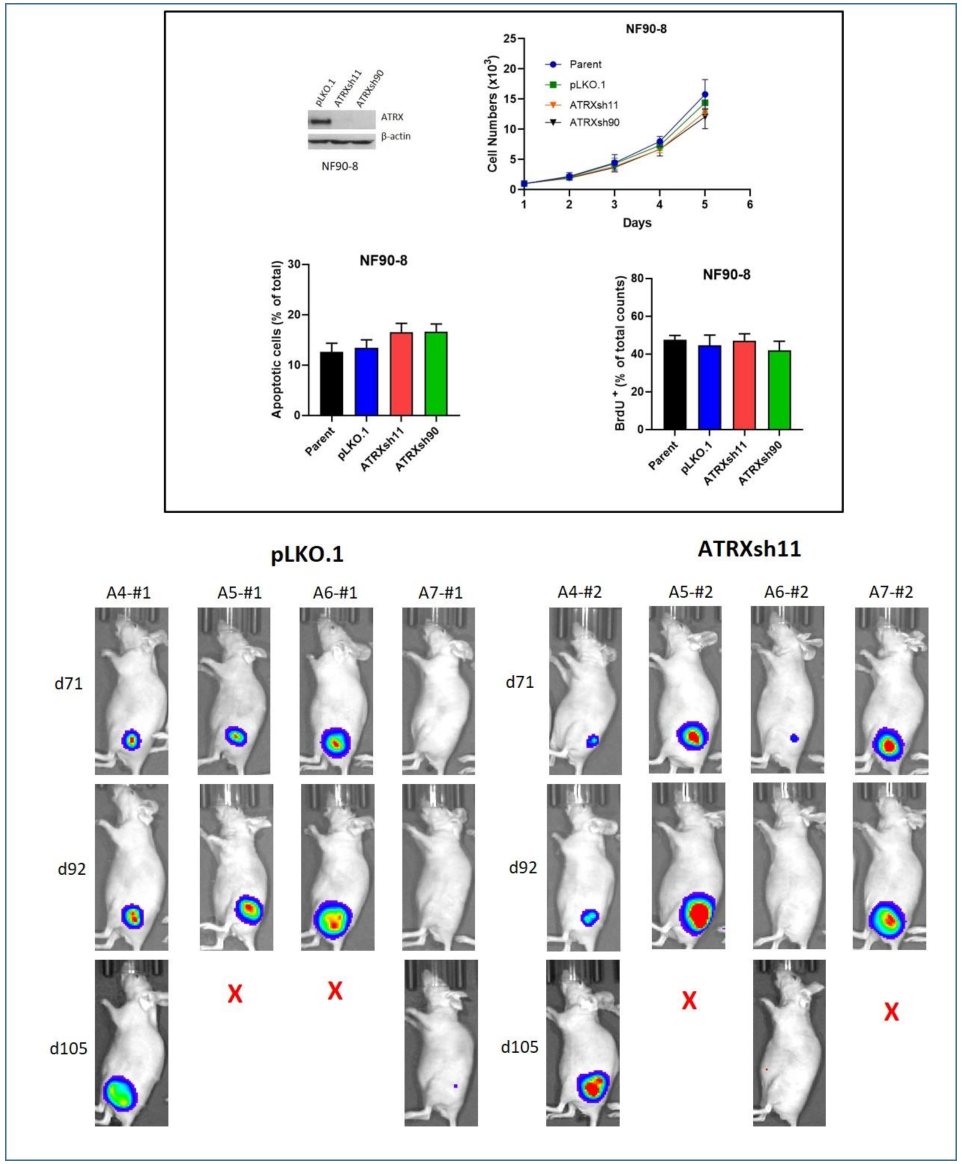

**Supplementary Figure 6. ATRX knockdown in ST88-14 has no effect on cell growth.** Successful ATRX knockdown in the MPNST cell line ST88-14 has no significant effect on cell growth *in vitro*.

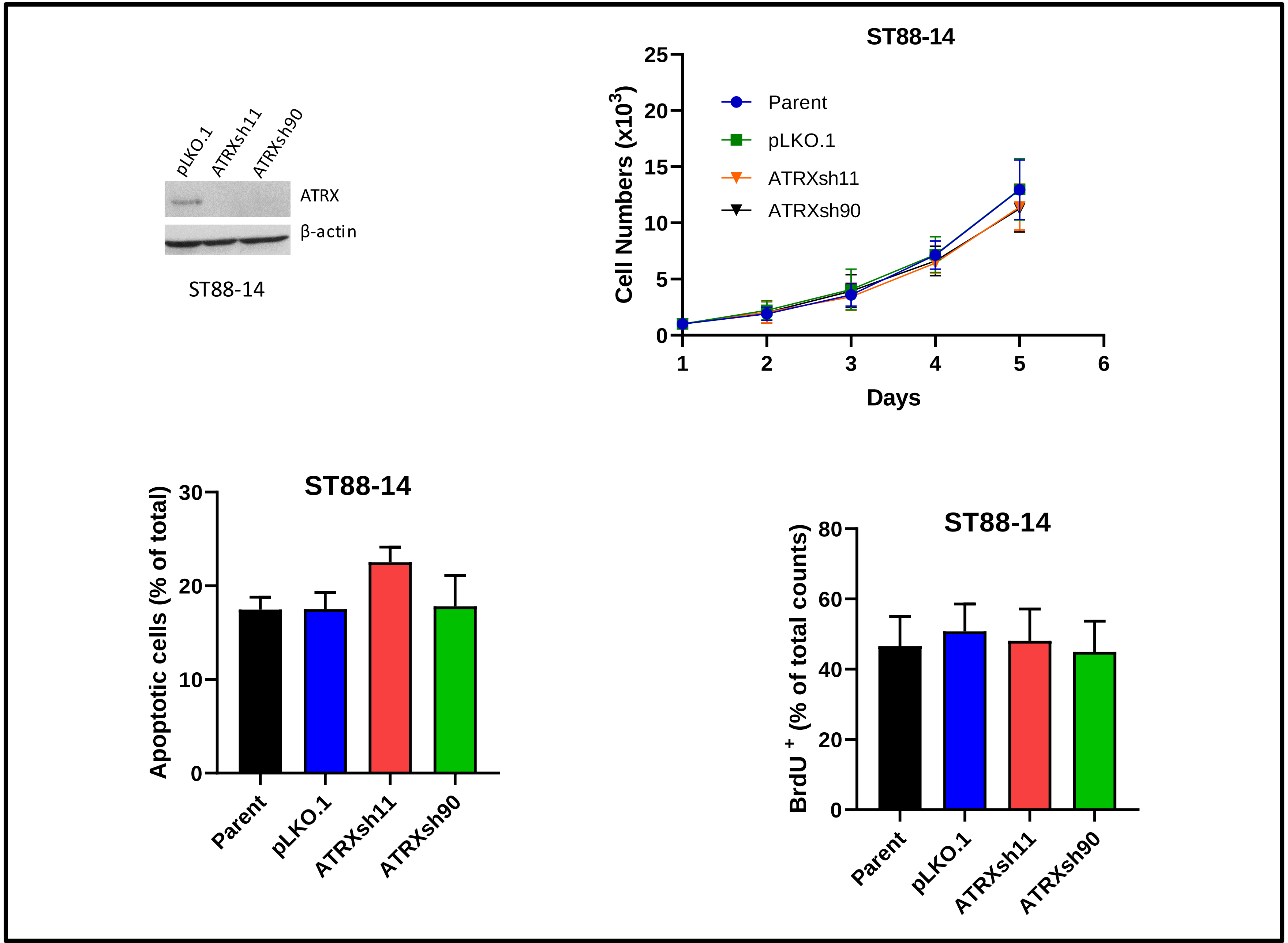

**Supplementary Figure 7. ATRX knockdown in NF1 sarcoma cell line has no effect on cell growth.** Successful ATRX knockdown in MPNST cell line JHH-CRC65 has no significant effect on cell growth *in vitro*. Significant decreases in ATRX levels but no effect on cell growth as demonstrated by the CellTiter-Blue assay (top) were detected. There was no increase in apoptosis or proliferation (BrdU+ cells, bottom).

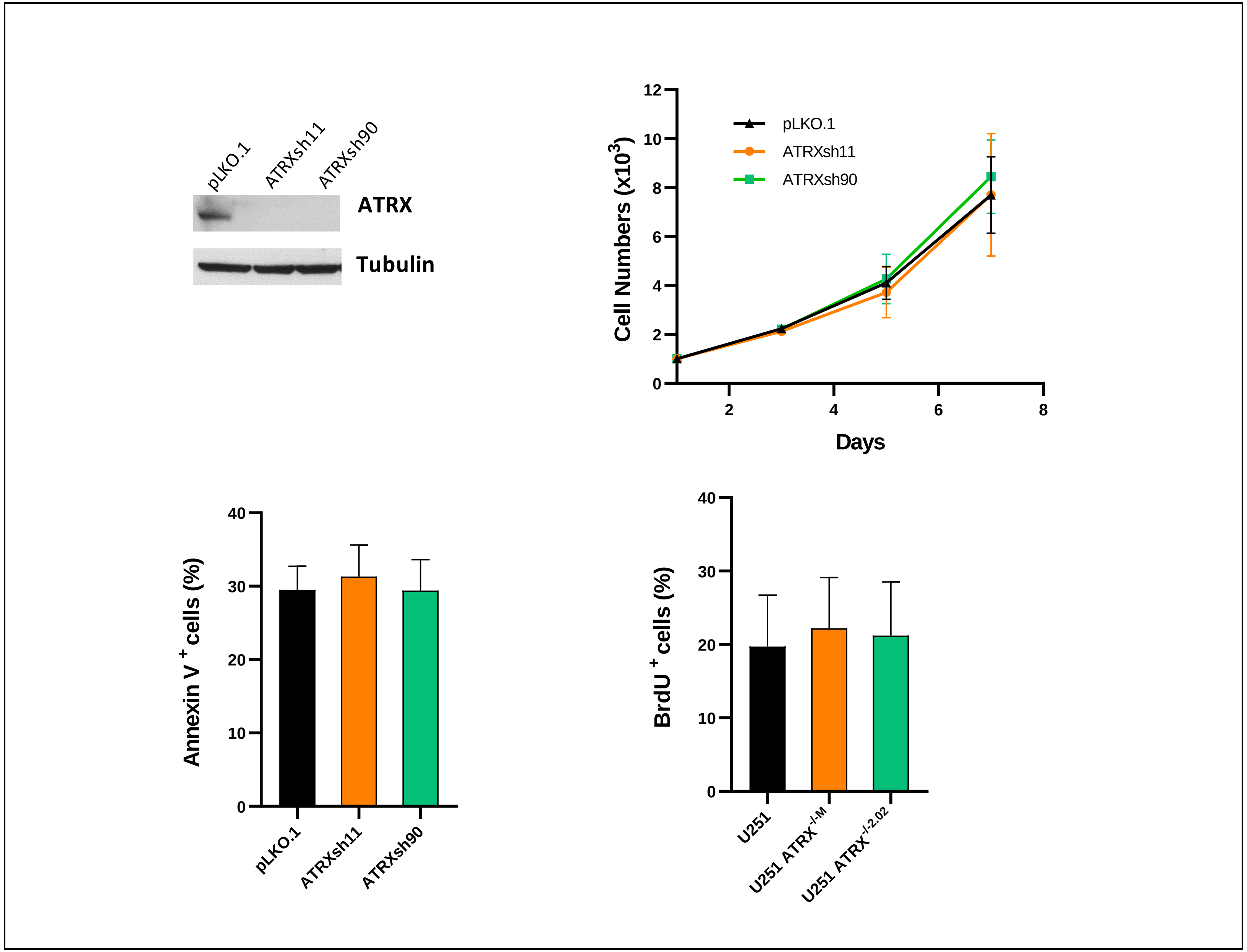

**Supplementary Figure 8. ATRX knockdown leads to senescence in non-neoplastic Schwann cells.** In contrast to MPNST cell lines, ATRX knockdown in a commercially available Schwann cell line resulted in senescence as demonstrated by increased mRNA levels of senescence marker CDKN1A.

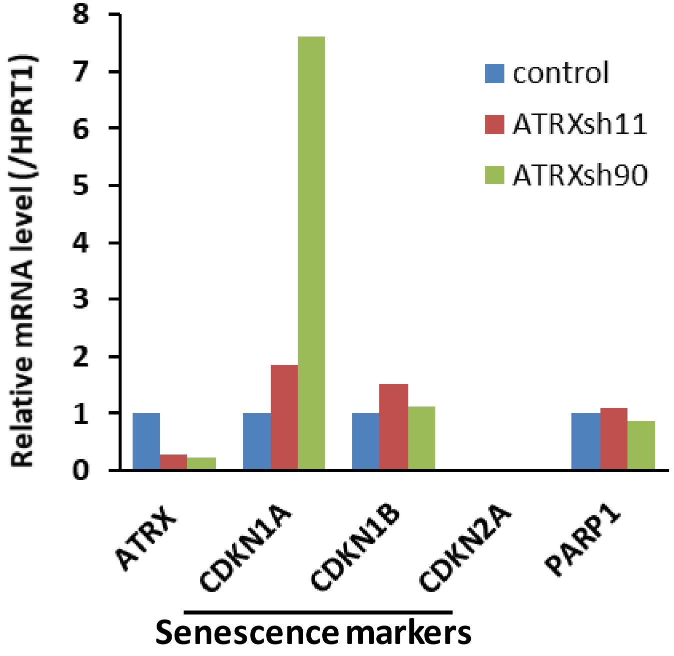

**Supplementary Figure 9 ATR inhibitor sensitizes U251 cells to temozolomide.** U251 cells were treated with various doses of AZD6738, temozolomide or combination for 5 days, cell survival were normalized with vehicle control (untreated).

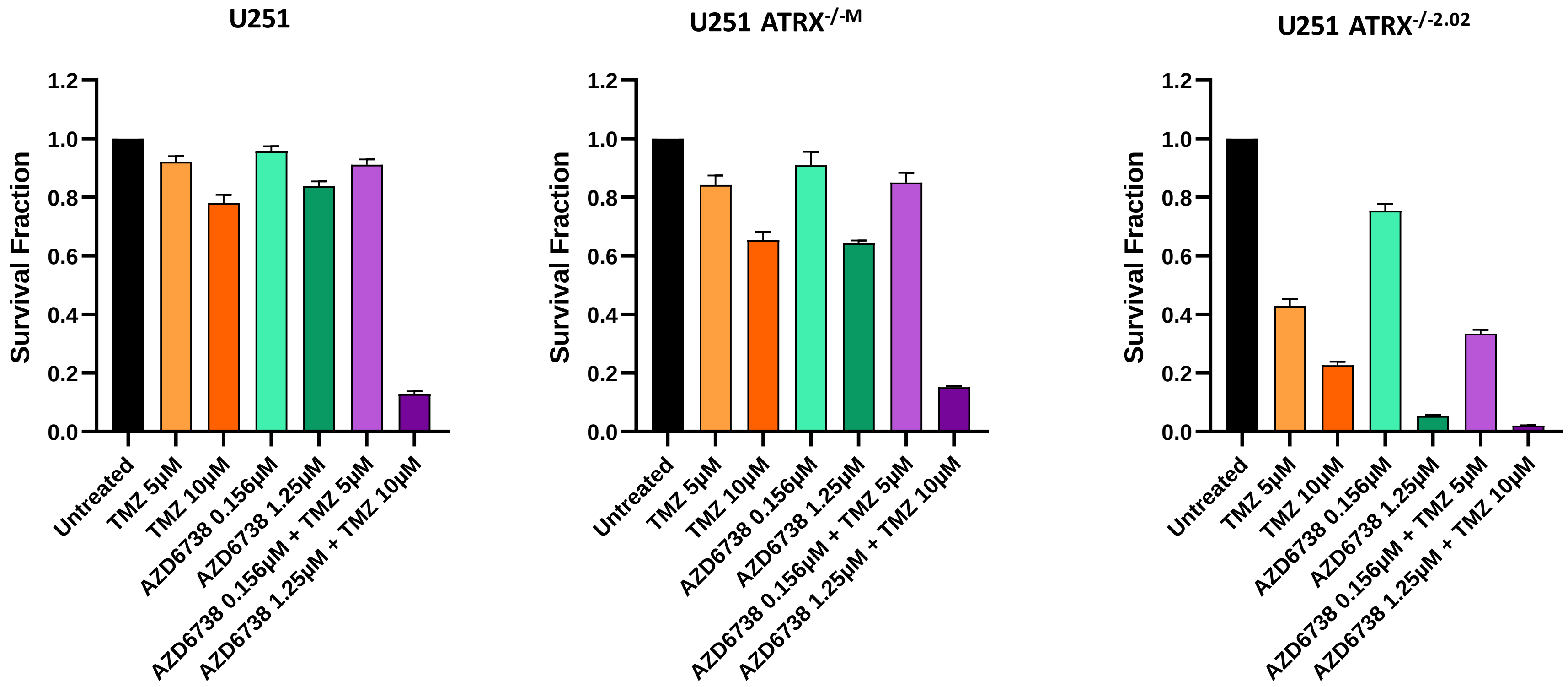

**Supplementary Figure 10. ATRX knockout sensitizes glioma cells to ATR inhibitor (AZD6738) but not MEK inhibitor (AZD6244).** MEK inhibitor AZD6244 results in decreased pERK, but not gH2AX levels (A). ATR inhibition did not sensitize U251 cells to MEK inhibition (AZD6244) (B).

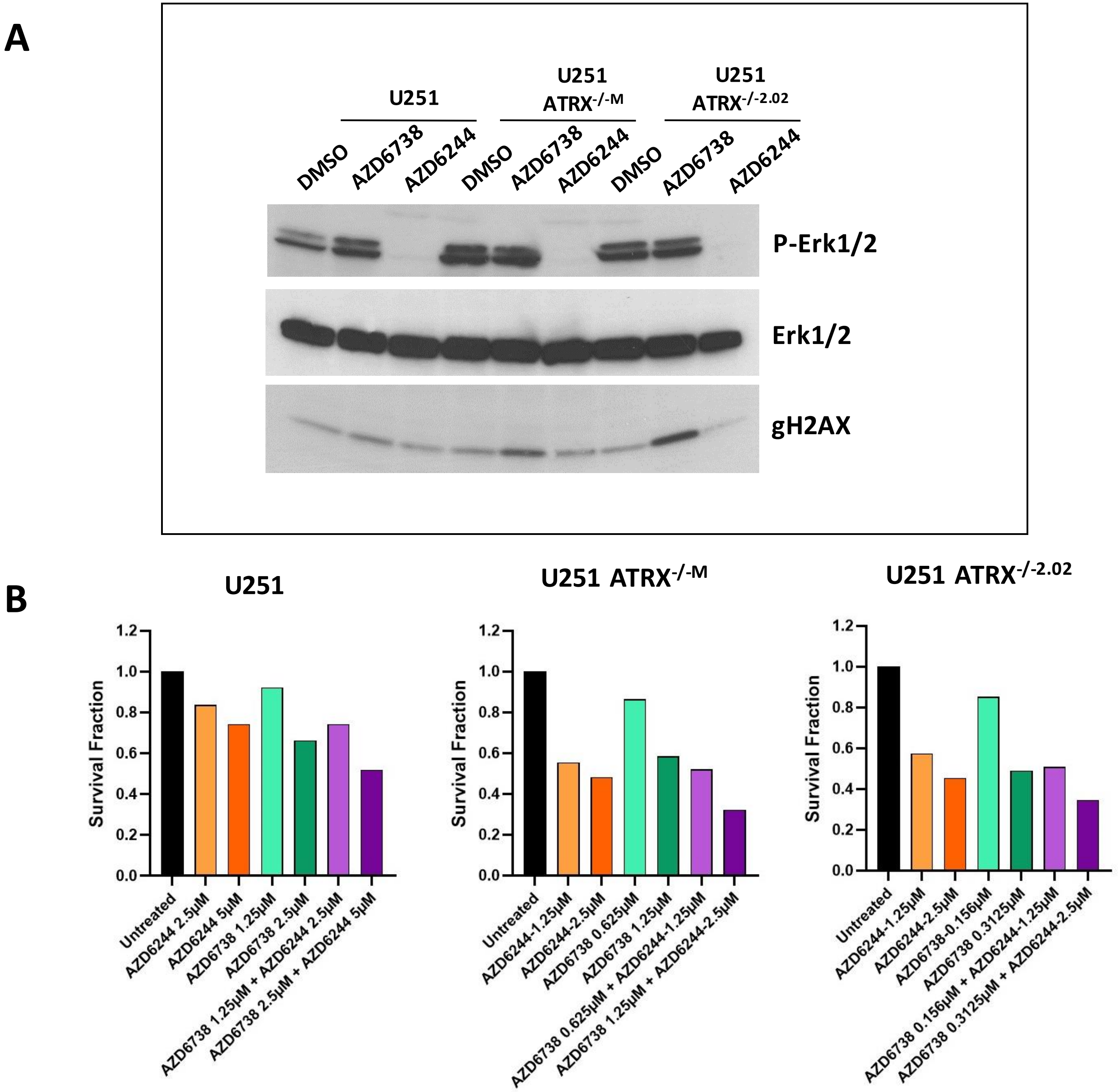

**Supplementary Figure 11. TERC knockout in NF90-8 inhibits telomerase activity.** TERC knockout was effective as demonstrated by pronounced decreased in TERC levels and telomerase activity.

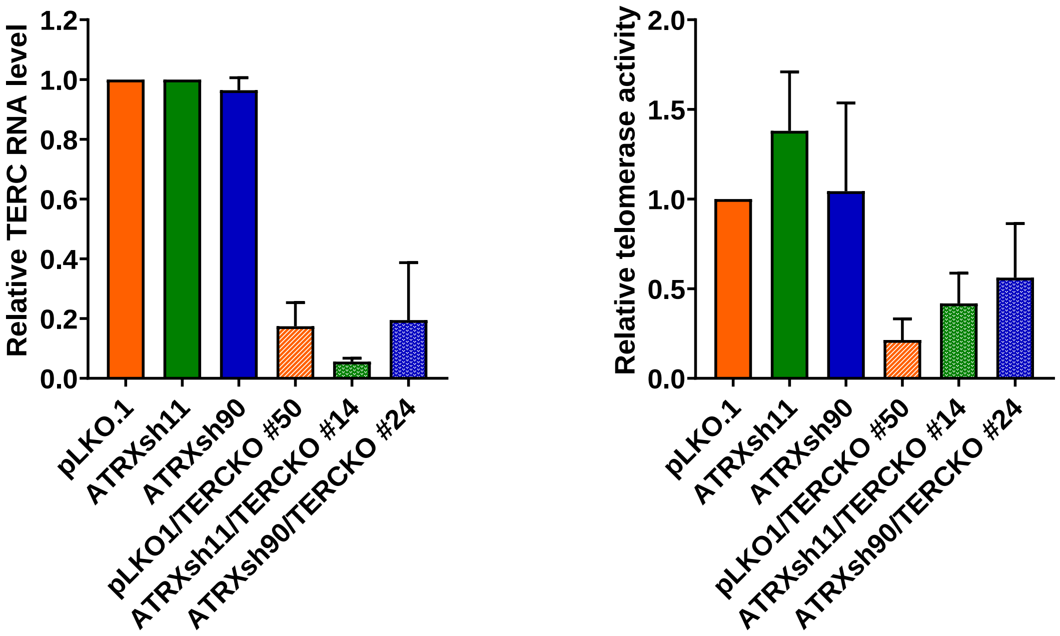

**Supplementary Figure 12. ATRX knockout has no effect on cell sensitivity to ATR inhibitor (AZD6738) and temozolomide.** No changes in cell growth of the NF1-derived sarcoma line JHH-CRC65 with and without ATRX knockdown after ATR inhibition or temozolomide as demonstrated by CellTiter Blue assay.

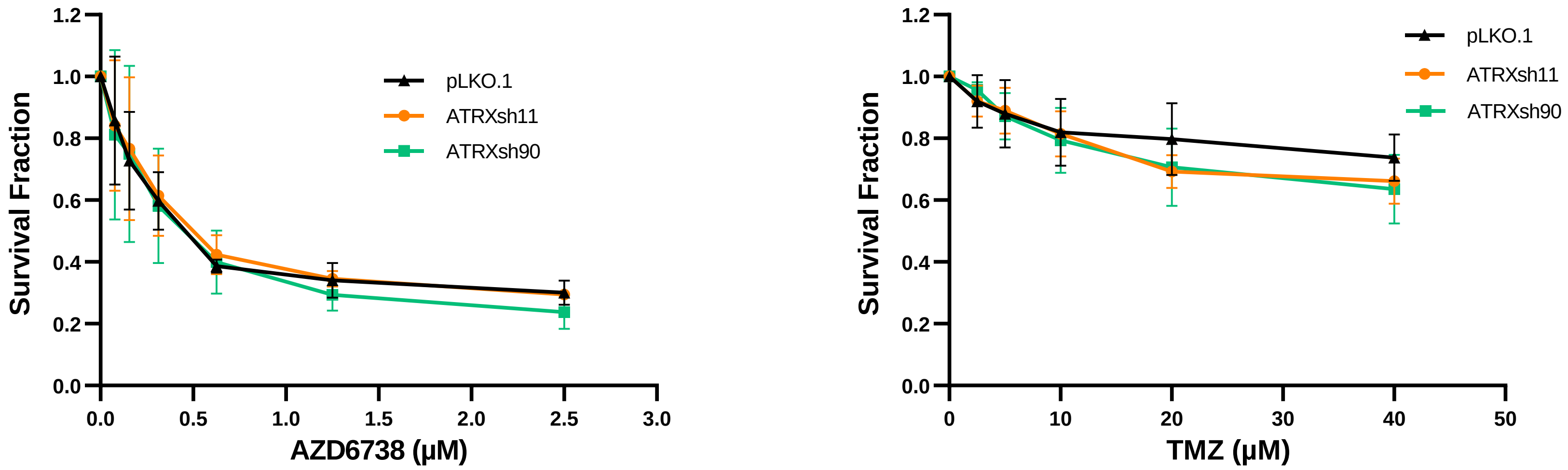
